# Estimating cellular pathways from an ensemble of heterogeneous data sources

**DOI:** 10.1101/006478

**Authors:** Alexander M. Franks, Florian Markowetz, Edoardo Airoldi

**Author notes:** This work was supported, in part, by NIH grant R01 GM-096193, NSF CAREER grant IIS-1149662, and by MURI award W911NF-11-1-0036 to Harvard University. EMA is an Alfred P. Sloan Research Fellow.

## Abstract

Building better models of cellular pathways is one of the major challenges of systems biology and functional genomics. There is a need for methods to build on established expert knowledge and reconcile it with results of high-throughput studies. Moreover, the available data sources are heterogeneous and need to be combined in a way specific for the part of the pathway in which they are most informative. Here, we present a compartment specific strategy to integrate edge, node and path data for the refinement of a network hypothesis. Specifically, we use a local-move Gibbs sampler for refining pathway hypotheses from a compendium of heterogeneous data sources, including novel methodology for integrating protein attributes. We demonstrate the utility of this approach in a case study of the pheromone response MAPK pathway in the yeast S. cerevisiae.

## 1 Introduction

Cellular mechanisms are driven by interactions between DNA, RNA, and proteins working together in cellular pathways. However, the current knowledge of information flow in the cell is still very incomplete (Kirouac et al., 2012). Even in well established signaling pathways studied for decades in model organisms, newer approaches can discover novel components (Möller et al., 2005) or cross-talk with other pathways (McClean et al., 2007). In cancer, finding pathways underlying disease development can lead to new drug targets (Balbin et al., 2013). This makes the dissection of cellular pathways one of the major challenges of systems biology and functional genomics.

One of the main obstacles to utilize high-throughput data in refining known pathway models is the gap between the relatively unbiased and hypothesis-free nature of generating genome-scale datasets and the need for very focused, hypothesis-driven research to test biological models in small or medium scale experiments (Hibbs et al., 2008). While researchers in computational biology usually start with a collection of data and reconstruct pathways from it, experimental biologists often start with a specific pathway hypothesis in mind and try to reconcile it with the evidence from high-throughput screens.

Here, we contribute to bridging this gap by introducing a comprehensive data integration strategy to refine a given pathway hypothesis. Our approach is characterized by three key features: First, we start with a *specific pathway model* and assess how well it is supported in a collection of complementary data sets. These data sets are heterogeneous and informative for distinct cellular locations. Second, we exploit this fact by introducing a *compartment specific* probabilistic model, where data types are only used for reconstructing the parts of a pathway they are informative about. Third, we explicitly include *node properties* in our model. This allows us to use data like protein phosphorylation states or protein domains, which have so far been underutilized for pathway structure learning (Ryan et al., 2013).

In this paper we show that our modeling approach can assist experimentalists in planning future studies by assessing which parts of a biological model are not well supported by data, and by proposing testable extensions and refinements of a given pathway hypotheses. We demonstrate the power of our approach in a case study in the yeast *S. cerevisiae.*

### Related work

Pathway reconstruction is a well established field in computational biology (Hyduke and Palsson, 2010; Markowetz and Spang, 2007). Several features distinguish our pathway refinement methodology from existing network reconstruction methods.

Comprehensive data integration strategies on large data collections were shown to be very successful in predicting protein function and interactions (Guan et al., 2012; Llewellyn and Eisenberg, 2008; Guan et al., 2008; Myers et al., 2005). These methods are very helpful for describing the global landscape of protein function, but offer less insight into individual molecular mechanisms and pathways. Our approach differs from methods to refine pathway hypotheses from expression profiles of down-stream regulated genes (Gat-Viks and Shamir, 2007), because we integrate heterogeneous data sources in a compartment-specific way.

We also differ from previous research on de-novo pathway reconstruction. These methods can be classified by how they use information about edges, paths and nodes in the pathway diagram for structure learning.

- Most approaches incorporate evidence for individual *edges* in the pathway diagram using phenotypic profiles (Mulder et al., 2012; Wang et al., 2012) or gene expression measurements (Li et al., 2013; Balbin et al., 2013; Schäfer and Strimmer, 2005a; Friedman, 2004; Segal et al., 2003), sometimes supplemented by additional data sources like transcription factor binding data (Bernard and Hartemink, 2005; Werhli and Husmeier, 2007) or protein-protein interactions (Gitter et al., 2013; Nariai et al., 2004; Segal et al., 2003). Other studies completely rely on protein-protein interactions to predict pathways (Mazza et al., 2013; Scott et al., 2006).
- Cause-effect relationships indicating *paths* from perturbed genes to observed effects are exploited in methods like SPINE (Ourfali et al., 2007), physical network models (Yeang et al., 2005), nested effects models (Wang et al., 2013; Markowetz et al., 2007; Tresch and Markowetz, 2008; Fröhlich et al., 2007, 2008) and others (Lo et al., 2012; Yip et al., 2010), with applications including DNA damage repair (Workman et al., 2006) and cancer signalling (Knapp and Kaderali, 2013; Stelniec-Klotz et al., 2012).
- *Node information*, i.e. features of individual proteins or genes, has been found useful for assigning proteins to pathways (Hahne et al., 2008; Fröhlich et al., 2008) but has so far been under-utilized in reconstructing pathway structure (Ryan et al., 2013).

Our method differs from de-novo pathway reconstruction in that we start with a hypothesis pathway and identify which hypothesized edges are supported by the data. We also differ from other methods which evaluate formal one and two sample network hypothesis tests (Yates and Mukhopadhyay, 2013). Our goal is not to explicitly to determine whether our initial hypothesis is “correct”- on the contrary we assume a priori that any initial hypothesis can be further refined and improved upon. In the spirit of FDR, we provide a list of edge probabilities that can assist experimentalists in their future studies. We assess which parts of an existing biological model are not well supported by a data as well as suggesting new edges which are supported by the data but which are not part of the original hypothesis. Further, we are the first to integrate data about *edges* and *paths* as well as *nodes* in the pathway diagram.

### Overview

We describe a compartment-specific probabilistic graphical model for posterior inference on cellular pathways in *section 2*, which can extend and refine a given biological model and predict novel parts of the pathway graph. Our model comprehensively integrates the three general types of data on edges, paths, and nodes. We demonstrate the utility of our methods in a case study in *S. Cerevisae (section 3)* by first exploring the information content in different data sources individually *(section 3.1)* and then evaluating results of posterior draws using both full data and leave-one-out data *(section 3.2).*

## 2 Integating high-dimensional responses of a cellular pathway

Given a set of a gene products, i.e., putative pathway members, we infer an undirected network model using a local-move Gibbs sampler. The pathway model, is defined in terms of N nodes and the edges between these pairs of nodes, *(n,m).* The edges are encoded by a binary random variable, *X*_*nm*_. The collection of edge-specific random variables defines the adjacency matrix, **X**, of the pathway model.

### Parameter estimation and posterior inference

The adjacency matrix **X** corresponding to the pathway model is latent since we cannot directly observe the edges. Thus, the primary goal of our analysis is to do posterior inference on the adjacency matrix, **X**, from a collection of M data sets, *Y*_1:*M*_. Although we treat **X** as latent, we differ from de-novo pathway reconstruction by incorportaing an informative hypothesis pathway which we use to train the models for data sets **Y**_1:*M*_ (see Section 3).

By Bayes rule, the posterior distribution on a pathway model,

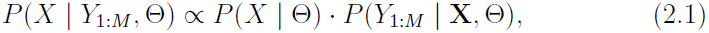

is proportional to the prior distribution on the pathway with the likelihood of the data. Here, Θ is a collection of prior parameters introduced below.

We use a local Gibbs sampling strategy to sample pathway models from posterior distribution in Equation 2.1. The sampler explores the space of pathway models by adding or removing edges in turn, one at a time. Specifically, the edge *X*_*nm*_ between gene products (*n*, *m*) is sampled according to a Bernoulli distribution, with probability of success

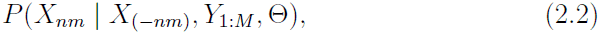

where *X*_(−*nm*)_ represents the set of edges without *X*_*nm*_.

#### 2.1 A compartment map defines context-specific data contributions

We use five complementary data types: physical binding of protein pairs (including yeast-two hybrid, mass spectrometry, and literature-curated data), transcription factor-DNA binding assays, gene knockout data, gene co-expression data, and node information (including protein domains and differential phosphorylation arrays) Importantly, different data sets can be very informative in specific cellular locations while completely uninformative in others. Thus, before we define the data likelihoods in section 2.2, it is essential to exploit this fact in our model. We translate expected compartment localization of a pair of gene products *(n,m)* into a binary importance vector 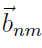 which drives the inference process by selecting the most informative data types for the compartments involved.

To instantiate the notion that different data are informative in different cellular locations, we introduce an additional modeling element: the compartment map, which contains three conceptual pathway compartments directly based on the organisation of the cell: First, the cell *membrane*, where receptor proteins sense signals from outside the cell; second, the *cytoplasm*, where protein cascades relay these signals to transcription factor proteins that enter the third compartment, the *nucleus*, to regulate the activity of target genes. The compartment map, *C*, is a 5 × 3 binary matrix that associates the three pathway compartments with the five data types to indicate which data type is informative about molecular interactions in which compartments (see Table 1).

**Table 1:** The compartment map, *C*, associates pathway compartments with those data types that are informative for such compartments. Prior information is informative for all compartments.

|  | Membrane | Cytoplasm | Nucleus |
| --- | --- | --- | --- |
| PPI | 1 | 1 | 0 |
| TF | 0 | 0 | 1 |
| Expr | 0 | 0 | 1 |
| Kout | 0 | 1 | 1 |
| Node | 0 | 1 | 0 |
| Prior | 1 | 1 | 1 |

In particular, each data set is described by a pair (*Y*_*i*_, *T*_*i*_), where *Y*_*i*_ denotes the collection of measurements, and *T*_*i*_ is five-level factor that denotes the specific data type used to collect the data and indexes the relevant row of *C*. We can now revise the form of the conditional distributions in Equation 2.2,

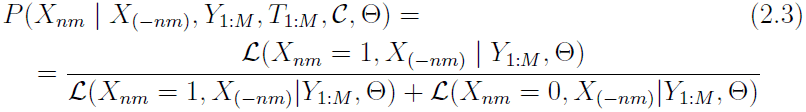

Overloading notation, we let *C*_*t*_(*n,m*) be an indicator reflecting whether the protein pair (*n,m*) is informative for data type t, based on the compartment map and the localizations of proteins n and m. This leads to the following likelihood specification:

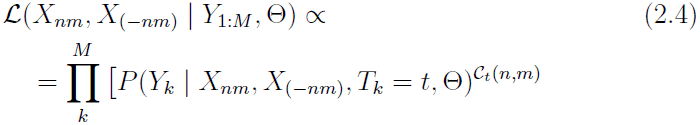

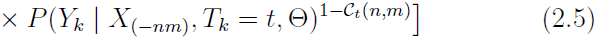

where the role of the indicator is to discard data collections from data types that are expected to carry little information about the protein pair of interest, according to information in *C*. That is, for any pair (*n,m*), *C*_*t*_(*n,m*) = 0 implies data set *Y*_*k*_ is conditionally independent of (*n, m*) given the rest of the pathway. In this case, the data in *Y*_*k*_ has no effect on the conditional posterior probability of *X*_*nm*_.

### 2.2 Modeling high-dimensional data for nodes, edges and paths

Data of different types need to be modeled differently. We focus on modeling five main data types: protein interaction data, protein-DNA binding data, gene co-expression data, gene perturbation data, and node attribute data (differential phosphorylation and protein domains). Below, we describe the likelihood functions corresponding to the main data types of interest.

#### Likelihood for protein interaction data

Here, we consider a single data set *Y*_*N×N*_ obtained with data type *T* aimed at measuring physical protein binding events (PPI). We reduce the likelihood of the data, *Y*, to a function the false positive and false negative rates, *α* and *β*. Given the pathway, *X*, we evaluate

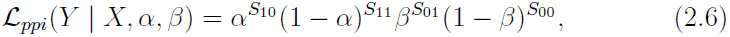

where *S*_*xy*_ counts the number of edges for which *X*_*nm*_ = *x* and *Y*_*nm*_ = *y.* For instance, *S*_10_ is the number of false positives.

#### Likelihood for protein-DNA binding data

Here, we consider a single data set *Y*_*N×K*_, obtained with data type *T* aimed at measuring transcription factor-DNA binding events (TF). Rather than hybridization levels (for ChIP-chip) or peaks (for ChIP-seq), we model the *p*-values corresponding to binding events, which makes our model independent of the technology used to detect the binding event. We develop a mixture model for the *p*-values, directly. Given the pathway, *X*, we expect to see a small p-value for protein *n* binding nucleotide sequence *m* whenever the edge *X*_*nm*_ is present. On the contrary, the *p*-values are uniformly distributed under the null hypothesis of no binding events, *X*_*nm*_ = 0. We evaluate

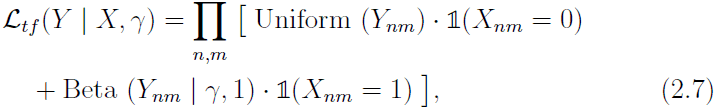

where 0 < *Y*_*nm*_ < 1 (*p*-value), and 0 < γ < 1. See a related beta-uniform mixture model introduced by Pounds and Morris (2003) in the context of multiple testing for differential expression.

#### Likelihood for knock-out data

Here we consider a data set *Y*_*M×N*_, where *Y*_*mn*_ is the log-two-fold change in expression of gene n, when gene m is knocked out. Let *Z*_*mn*_ be a binary variable representing the existence of a directed path from gene n to gene m, *through a transcription factor.* While we consider the set of undirected pathway models, we temporarily impute directionality using the fact that the cellular signal should flow from the cytoplasm to the nucleus. We model the knockout data as a mixture of normals:

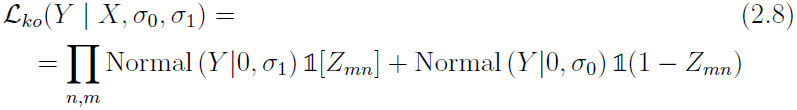

The standard deviations for change in expression are represented by σ_0_ (when there is no path between the knockout and a target) and σ_1_ (there is a path). The assumption is that σ_1_ > σ_0_ since we expect a larger change in expression of n for knockout m when n and m are connected in the pathway.

#### Likelihood for gene co-expression data

Here, we consider a single data set *Y*_*N×N*_ aimed at measuring gene expression. Rather than hybridization levels (for microarrays) or the number of reads (for mRNA sequencing), we model correlations among the profiles of pairs of genes, which again makes our model independent of the details of the measurement technology. We develop a mixture model for the correlations, directly. Given the pathway, *X*, we expect to see correlation between the expression profiles of two genes whenever they are co-regulated. Similarly to Schäfer and Strimmer (2005b), we use a mixture model for the distribution of the sample correlation coefficient 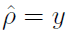 of the form

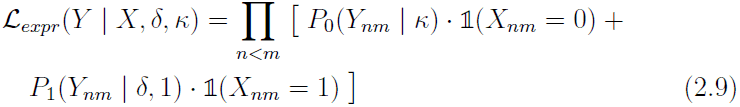

When *X*_*nm*_ = 0, we expect the two gene profiles to be uncorrelated. Differently from Schäfer and Strimmer (2005b), however, we chose a distribution that puts more emphasis on higher correlation if we see an edge in the model, *X_nm_* = 1, using a one-parameter beta distribution,

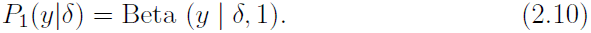

#### Likelihood for node attributes data

Here, we consider a single data set *Y*_*M×N*_ that lists node-specific attributes such as protein domains from PFAM (Punta et al., 2012) and SMART (Schultz et al., 1998; Letunic et al., 2012) databases, and differential phosphorylation data (Gruhler et al., 2005). We develop novel techniques to model protein attributes. Specifically, we model the likelihood of an attribute conditional on the given pathway **X**. We term our models for node attributes “relation regression.” For differential phosphorylation data, *Y*_*N×1*_,

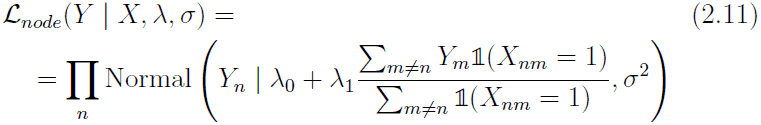

In other words, the differential phosphorolation, *Y_n_*, is assumed to be linearly related to the mean differential phosphorolation of the neighbors of node n. Similarly, for the protein domain data, *D*_*N×K*_, we use an auto-logistic regression to model the data. Specifically, for *D*_*nk*_, a binary variable indicating the presence of domain k in protein n,

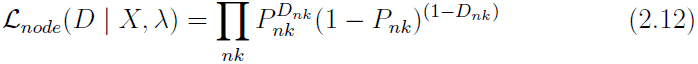

where

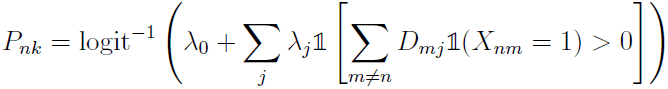

Here, logit(*P*_*nk*_) is linearly related to the presence of domains in neighboring genes. In both the normal and logistic regression cases, we fit the regression coefficients, 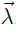, using our initial pathway hypothesis. In the logistic model, we use a weakly-informative Cauchy prior for the coefficients (Gelman, 2008). This controls for any overfitting and separation problems that may occur.

#### Prior distribution on the space of pathway models

In this study our focus lies on assessing the extent to which the data support a pathway model *X*. We choose a block model prior *P*(*X*) over binary matrices of size *N* × *N* with edge density fixed by compartment. In general, any informative prior distribution on graphs could be used here to encode biological knowledge (Isci et al., 2013; Mukherjee and Speed, 2008).

## 3 Analysis of the pheromone response pathway in *S. cerevisiae*

To demonstrate the efficacy of our approach, we examine the pheromone response MAPK pathway in the yeast *S. cerevisiae.* It offers the opportunity to combine a large collection of datasets with a solid understanding of the pathway structure. The pheromone pathway is the subject of intense research efforts in computational biology as well as experimental biology (Hara et al., 2012; Scott et al., 2006; Kofahl and Klipp, 2004) and shows cross-talk to other MAPK pathways (Nagiec and Dohlman, 2012; McClean et al., 2007; Gat-Viks and Shamir, 2007).

### Initial pathway construction

To start our analysis in a way relevant to refining and extending existing knowledge of signaling pathways, we extracted a model of the pheromone response pathway from the summary of MAPK pathways (sce04010) in the database KEGG (Kanehisa and Goto, 2000) and combined it with known transcription factor (TF) targets from two independent studies (Simon et al., 2001; Ren et al., 2000).

We split the pathway into three parts: the *membrane* compartment containing the receptor proteins, the *cytoplasm* compartment containing the MAPK cascade to activate the transcription factors (TF), and the *nuclear* compartment containing the TFs and their targets. Figure 1-A depicts the pathway hypothesis. Proteins mediating between two compartments (like TFs) are contained in two sub-graphs and marked by grey boxes. TF targets that are also members of other compartments are indicated in bold.

**Figure 1:**
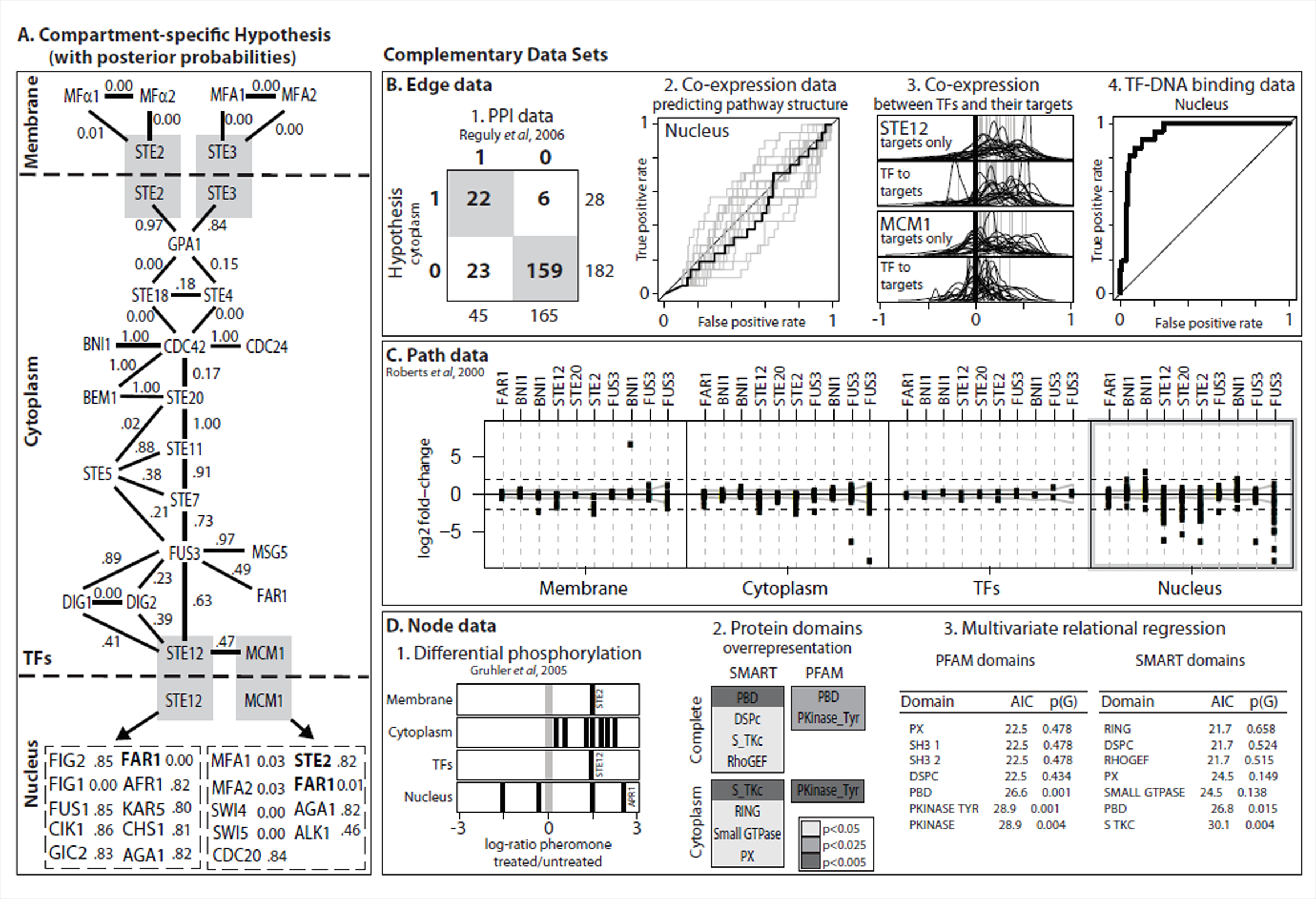
Compartment-specific pathway hypothesis, posterior probabilities, and evaluation of support in the data. **A.** Pathway hypothesis and posterior edge probabilities for the Yeast pheromone response pathway. The numbers by each edge reflect the “posterior probability”. **B.** Edge data: (1) protein- protein interactions in the cytoplasm, (2) gene co-expression in the nucleus, (3) co-expression of TFs with their targets and between targets for STE12 and MCM1, and (4) TF binding data. **C.** Cause-effect data. D.Node data: (1) Differential phosphorylation, (2) Overrepresentation of protein-domains in different compartments, (3) goodness-of-ht of auto-logistic models on protein domains from PFAM and SMART.

### 3.1 Exploratory data analysis of individual data types

Before inferring the full model from all data, we explored the information content in each type of data individually.

#### Protein-protein interactions (PPI)

We compared data from several complementary high-throughput assays, all available from BioGRID (Stark et al., 2006) as well as a literature-curated dataset (Reguly et al., 2006). We analyzed the overlap between the protein interactions and the pathway hypothesis of Fig 1-A. None of the datasets are informative for the membrane and nuclear compartments. Surprisingly, in the cytoplasm compartment we found that all of the high-throughput datasets show only ≤ 3 interactions between any of the proteins in the pathway. The situation was very different for the literature-curated data. Here, 45 interactions in the cytoplasm compartment covered 22 out of the 28 edges there (sensitivity > 78%, specificity > 87%, see Fig 1-B1).

#### TF-DNA binding data

We used the transcription factor binding data of (Harbison et al., 2004), which is independent of our definition of TF targets in the pathways hypothesis. The ROC in Figure 1-B4 shows very clear signal to distinguish the targets posited in the biological model from all other pathway genes.

#### Co-expression data

For gene expression data, we examined datasets in which the pathway genes showed a significant difference in correlation structure from all other yeast genes (using the SPELL algorithm of (Hibbs et al., 2007)) resulting in 20 datasets from 15 publications (including Roberts et al., 2000; Gasch et al., 2000; Brem and Kruglyak, 2005). Figure 1-B2 shows ROCs for predicting edges in the nuclear compartment for all datasets (grey lines) and the concatenated data (black line). No curve improves much on random prediction (the main diagonal). The reason is biological: Because expression data are a poor surogate for protein activity, TFs are often less well correlated to their targets than the targets are between each other (Figure 1-B3). For STE12, which regulates itself, all correlation coefficients exhibit a strong trend towards high positive correlation. Whereas MCM1, which is not self-regulating, is far less strongly correlated to its targets than the targets are between each other. Thus, in general it is more informative to use the correlation between targets for inference, which is consistently high whether or not a TF is transcriptionally regulated itself.

#### Gene perturbation data

Paths in the graph are visible in cause-effect datasets (Hughes et al., 2000; Roberts et al., 2000). We find only very small effects of perturbations in the pathway on the expression of members of the membrane and cytoplasm compartment including TFs. Figure 1-C summarizes this result for the Roberts et al. (2000) data. Very similar results were found for the Hughes et al. (2000) data. The four boxes correspond to the three compartments plus TFs. In each box, a vertical line corresponds to a perturbation in the pathway (some replicated). The dots show the fold-changes of the pathway genes in this compartment. Only in the nuclear compartment are wide-spread large fold-changes visible. This observation motivates the construction of our likelihood around the presence of paths between the knockout and genes in the nuclear compartment (see section 2).

In this way, when the knockout is far enough upstream, there is information about edges in the cytoplasm as well, even if the proteins there show no effect on the transcriptional level.

#### Protein phosphorylation

A first example of node information is protein phosphorylation. The study of Gruhler et al. (2005) assessed differential phosphorylation of proteins in response to pheromone. Figure 1-D1 shows the log-ratios between the pheromone treated and untreated conditions. Almost all proteins of the pheromone pathway measured by Gruhler et al. (2005) are up-regulated, which makes sense for a kinase cascade. The phosphorylation we observe for proteins corresponding to genes only attributed to the nuclear compartment in our model must be due to other kinase pathways in the cell. We further assessed to what extent the differential phosphorylation is correlated with the pathway model by fitting an auto-logistic regression. As a measure of correlation we computed the variance explained, *R*_2_ = 0.76, using the bootstrap. The variance explained by the auto-logistic regression was found statistically significant, when compared to the correlation of differential phosphorylation with randomized pathway models, *p* ≈ 0.062, and with randomized protein permutations on the true pathway model, *p* ≈ 0.059.

#### Protein domains

A second example of node information are protein domains. We retrieved protein domains from PFAM (Punta et al., 2012) and SMART (Letunic et al., 2012). First, we sought to quantify which domains, if any, were over-represented in the set of proteins involved in the complete pheromone response pathway as well as in each compartment, in turn. Figure 1-D2 lists the domains that were found to be over-represented in the complete pathway and in the cytoplasm; darker shades of gray indicate a more significant p-value for the over-representation test.

Second, we sought to quantify to what extent the presence or absence of specific protein domains in proteins interacting with a given protein, *P*, was informative about the presence or absence of the same domain in such protein, *P*. This analysis was carried out using auto-logistic models, which summarize the informativeness of protein domains between interacting proteins on average, across all proteins in a given pathway. We fit auto-logistic regressions using each protein *P* in the cytoplasm compartment of the pheromone response pathway as data point, and the presence or absence of domains *D*_*1:K*_ in any one protein among those interacting with *P* as covariates.

We fit multivariate models, which assume that the presence or absence of either the same or complementary domains is a factor that facilitates protein physical interactions. The two tables in 1-D3 summarize the goodness of fit of the multivariate models, and report bootstrap p-values to assess the significance of the AIC scores. Figure 1-D3 shows the p-values obtained by fitting the multivariate auto-logistic regression to randomized pathway models. The domains identified by the multivariate models as putatively carrying signal about the pheromone pathway in the cytoplasm overlap with the domains identified by the over-representation analysis above; namely, P21 rho-binding domains, S-TKc domains, and tyrosine-specific catalytic domains.

In summary, node attributes of the proteins involved in the pheromone response pathways are informative about mechanistic elements of the kinase cascade, across cellular localizations and in the cytoplasm. These findings suggest that integrating node attributes such as protein domains and cellular localization should increase the likelihood of pathway models that encode real biological signal about the inner working of a target pathway.

#### Data Integration

The previous results suggest that some datasets are indeed more informative in certain cellular locations. For example, protein interactions can explain wide parts of the kinase cascade in the cytoplasm, while co-expression is very strong for TF targets. However, no dataset is informative in all compartments: Neither protein interactions nor knockout data can explain a complete pathway. The pheromone response pathway is an archetypical MAPK pathway, so we expect these observations also to be valid for other MAPK and signaling pathways. These results suggest that the compartment-specific modeling approach we take here is sensible. As a proof of concept, we use the results of exploratory data analysis to heuristically construct the compartment map, *C* (Table 1). Ultimately, we hope to infer the compartment map in a statistically principled way.

### 3.2 Validation of the integrated analysis

We evaluated how well the joint model, which combines all the complementary data types discussed above, supports the pathway hypothesis in Section 3 by sampling 1000 possible pathways using MCMC and tabulating the posterior probabilities over the edges.

The logistic regression model for domain data may be subject to overfitting and separation. This can occur since there are many different protein domains present, yet the frequency of any single domain is fairly low. To mitigate this issue, we used a Cauchy prior on the coefficients for the suto-logistic regression, which is a sensible default prior for this model (Gelman, 2008). Since the domain information in the pheromone pathway is relatively sparse, we also collected protein domain data from other MAPK pathways and used the hypothesized structure of those pathways to help learn the regression coefficients. Figure 1A includes the posterior probabilities for the edges in our initial hypothesis.

We also used a *leave-one-out* strategy to evaluate the predictive power of our model. We ran 37 separate simulations where each node was in turn left out of the training pathway. The edges connected to this node were propagated to the neighboring nodes of the left-out node. We left out the nodes rather than edges, because specifically leaving out edges is equivalent to assuming that we know there is no edge present. We needed to construct our model in a way that encodes ignorance about the presence of an edge. Leaving out the nodes, instead of the edges, is one way of being agnostic about the presence of edges attached to that node. Only the coefficients in the auto-logistic regression were learned from the pathway hypothesis, so only the node likelihoods were affected. Table 2 shows the posterior probabilities for edges (under simulations in which a node was removed from the prior hypothesis pathway). This table presents posterior probabilities for edges involved in knockout experiments.

**Table 2:**
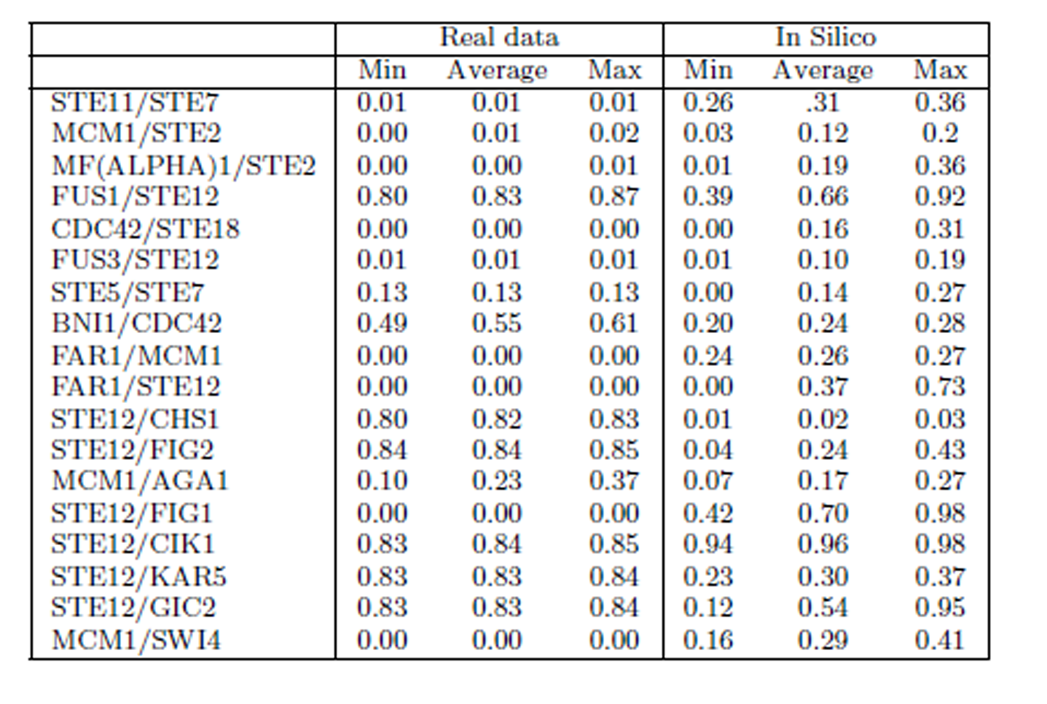
Posterior edge probabilities for leave-one-out trials involving edges in knockout experiments. Since we use a leave-node-out scheme, there are two posterior probabilities for an edge (corresponding to which of the two node endpoints were left out for that particular simulation).

Lastly, Figure 2 shows the precision-recall curve for our model, by compartment. For the membrane compartment, only the PPI data is informative, and weakly so. Thus, it performs the most poorly, although there are also by far the fewest genes in this compartment. By contrast, the nuclear and cytoplasm compartments both have high precision and recall.

**Figure 2:**
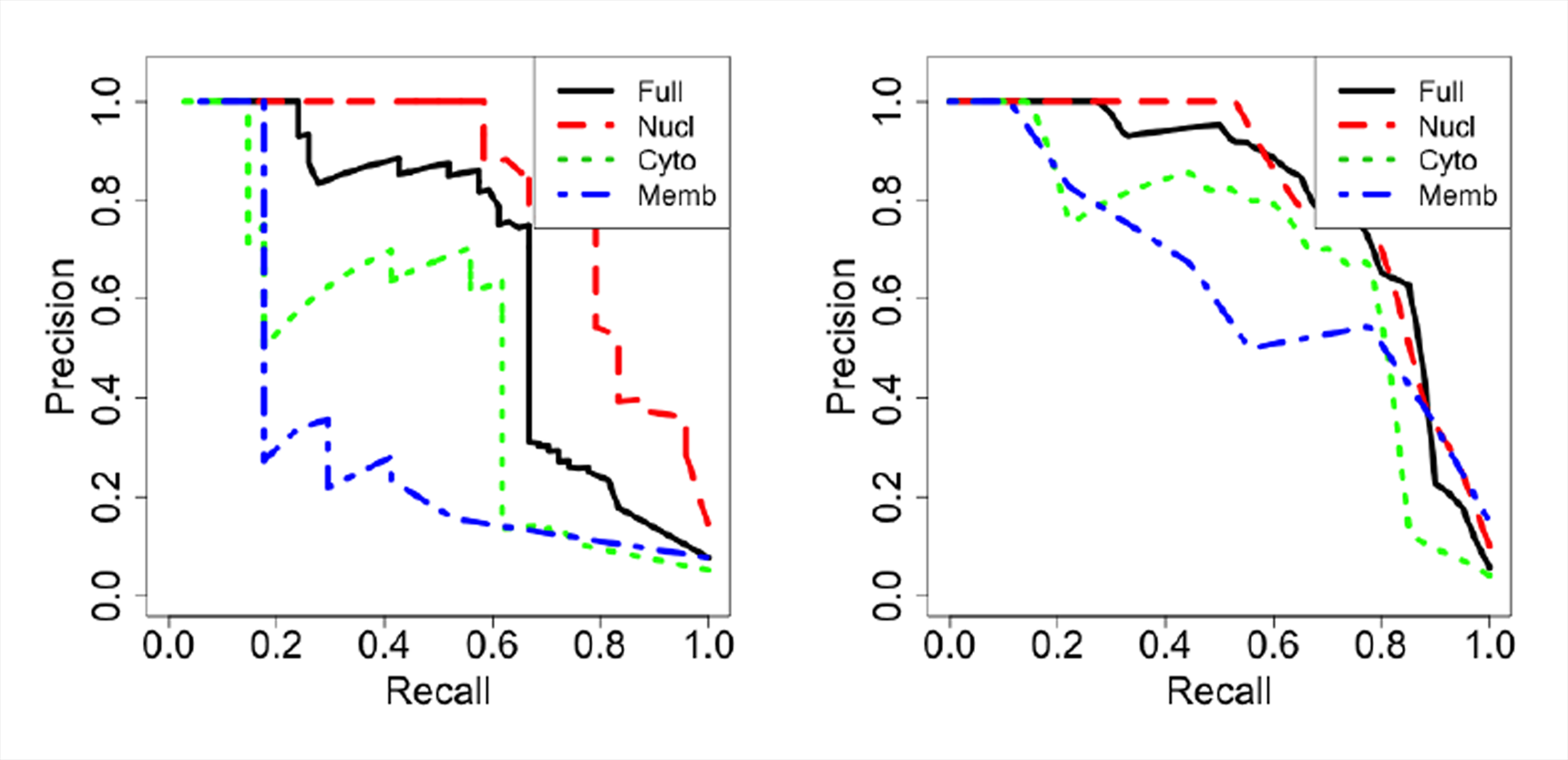
Precision/Recall curves overall and by compartment for the MAPK pathway (left) and simulated data (right). In truth, the membrane compartment, which has the fewest genes, performs poorly because only the PPI dataset is (weakly) informative there. The simulated data curve reflects the average Precision/Recall over 30 simulated datasets.

### 3.3 Inferring cross-talk with other pathways

With our model, we are also able to identify possible cross-talk between pathways. In this paper, we focus on the pheromone response pathway, but our model can easily be used on other pathways, as long as we specify the relevant genes and transcription factors, and their corresponding cellular locations.

For instance, the MAPK pathway consists of the pheromone sub-pathway, as well as hypotonic shock, osmolarity and starvation sub-pathways. The degree of interaction between components of these MAPK pathways is not currently known. To identify cross-talk between the pheromone pathway and other MAPK pathways, we can simply include a new set of genes from the other sub pathways and fit the model as usual. The results for the cross-talk evaluations are displayed in Table 3.

**Table 3:**
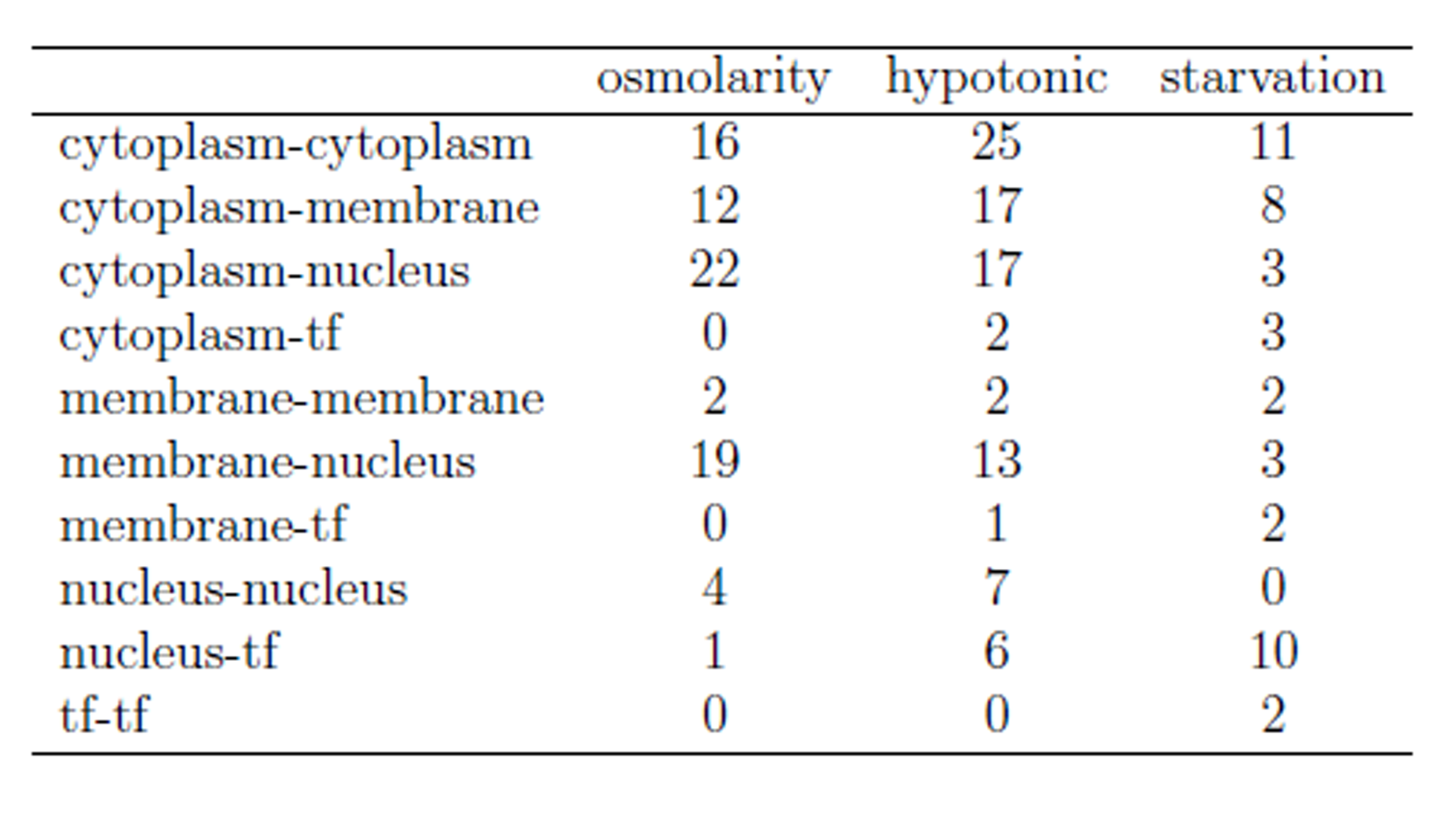
Number of inferred edges between the pheromone pathway and one of the other three sub-pathways with posterior probabilities above 0.3.

**Table 4:** Posterior edge probabilities.

|  | Gene 1 | Gene 2 | Prob |
| --- | --- | --- | --- |
| 1 | STE12 | DIG2 | 0.60 |
| 2 | STE12 | FUS1 | 0.39 |
| 3 | STE12 | FUS3 | 0.01 |
| 4 | STE12 | FAR1 | 0.00 |
| 5 | STE12 | MCM1 | 0.00 |
| 6 | STE12 | FIG2 | 0.43 |
| 7 | STE12 | FIG1 | 0.42 |
| 8 | STE12 | CIK1 | 0.98 |
| 9 | STE12 | GIC2 | 0.12 |
| 10 | STE12 | AFR1 | 0.01 |
| 11 | STE12 | KAR5 | 0.23 |
| 12 | STE12 | CHS1 | 0.03 |
| 13 | STE12 | AGA1 | 0.27 |
| 14 | DIG2 | STE12 | 0.68 |
| 15 | DIG2 | FUS3 | 0.00 |
| 16 | STE7 | STE11 | 0.26 |
| 17 | STE7 | STE5 | 0.21 |
| 18 | STE7 | FUS3 | 0.26 |
| 19 | STE11 | STE7 | 0.36 |
| 20 | STE11 | STE20 | 0.24 |
| 21 | STE11 | STE5 | 0.00 |
| 22 | STE20 | STE11 | 0.00 |
| 23 | STE20 | CDC42 | 0.31 |
| 24 | STE20 | BEM1 | 0.08 |
| 25 | STE20 | STE5 | 0.00 |
| 26 | CDC42 | STE20 | 0.00 |
| 27 | CDC42 | BNI1 | 0.28 |
| 28 | CDC42 | STE4 | 0.24 |
| 29 | CDC42 | STE18 | 0.31 |
| 30 | CDC42 | BEM1 | 0.47 |
| 31 | CDC42 | CDC24 | 0.50 |
| 32 | FUS1 | STE12 | 0.98 |
| 33 | BNI1 | CDC42 | 0.20 |
| 34 | MFA1 | STE3 | 0.34 |
| 35 | MFA1 | MCM1 | 0.07 |
| 36 | STE2 | MF(ALPHA)2 | 0.01 |
| 37 | STE2 | GPA1 | 0.35 |
| 38 | STE2 | MCM1 | 0.20 |
| 39 | STE3 | MFA1 | 0.30 |
| 40 | STE3 | GPA1 | 0.13 |
| 41 | MF(ALPHA)2 | STE2 | 0.36 |
| 42 | GPA1 | STE2 | 0.01 |
| 43 | GPA1 | STE3 | 0.14 |
| 44 | GPA1 | STE4 | 0.14 |
| 45 | GPA1 | STE18 | 0.12 |
| 46 | STE4 | CDC42 | 0.22 |
| 47 | STE4 | GPA1 | 0.14 |
| 48 | STE18 | CDC42 | 0.00 |
| 49 | STE18 | GPA1 | 0.13 |
| 50 | BEM1 | STE20 | 0.36 |
| 51 | BEM1 | CDC42 | 0.18 |
| 52 | CDC24 | CDC42 | 0.18 |
| 53 | STE5 | STE7 | 0.00 |
| 54 | STE5 | STE11 | 0.00 |
| 55 | STE5 | STE20 | 0.00 |
| 56 | STE5 | FUS3 | 0.00 |
| 57 | FUS3 | STE12 | 0.19 |
| 58 | FUS3 | DIG2 | 0.21 |
| 59 | FUS3 | STE7 | 0.22 |
| 60 | FUS3 | STE5 | 0.05 |
| 61 | FUS3 | MSG5 | 0.05 |
| 62 | FUS3 | FAR1 | 0.00 |
| 63 | MSG5 | FUS3 | 0.00 |
| 64 | FAR1 | STE12 | 0.73 |
| 65 | FAR1 | FUS3 | 0.27 |
| 66 | FAR1 | MCM1 | 0.27 |
| 67 | MCM1 | STE12 | 0.00 |
| 68 | MCM1 | MFA1 | 0.15 |
| 69 | MCM1 | STE2 | 0.03 |
| 70 | MCM1 | FAR1 | 0.24 |
| 71 | MCM1 | SWI4 | 0.41 |
| 72 | MCM1 | MFA2 | 0.20 |
| 73 | MCM1 | AGA1 | 0.27 |
| 74 | MCM1 | ALK1 | 0.15 |
| 75 | MCM1 | SWI5 | 0.38 |
| 76 | MCM1 | CDC20 | 0.34 |
| 77 | SWI4 | MCM1 | 0.16 |
| 78 | MFA2 | MCM1 | 0.19 |
| 79 | FIG2 | STE12 | 0.04 |
| 80 | FIG1 | STE12 | 0.98 |
| 81 | CIK1 | STE12 | 0.94 |
| 82 | GIC2 | STE12 | 0.95 |
| 83 | AFR1 | STE12 | 0.02 |
| 84 | KAR5 | STE12 | 0.37 |
| 85 | CHS1 | STE12 | 0.01 |
| 86 | AGA1 | STE12 | 0.00 |
| 87 | AGA1 | MCM1 | 0.07 |
| 88 | ALK1 | MCM1 | 0.24 |
| 89 | SWI5 | MCM1 | 0.13 |
| 90 | CDC20 | MCM1 | 0.18 |

### 3.4 Performance assessment on simulated data

We also fit the model to *in silica* data. We constructed the “true pathway” to match the hypothesized MAPK pheromone pathway of Figure 1A. That is, we fixed a pathway with the matching nodes and edges. We then generated in silico datasets from the models specified in Section 2. The one exception is the data generation for the node data.

Here, we generate the presence of domains in a way such that short chains in the pathway are more likely to share domains than are random non-neighboring nodes. Specifically, we randomly chose chains of length 1 to 4 and added a common “domain” to every node in that chain. In this way, the domain data realistically reflect the notion that genes sharing common protein domains are more likely to interact.

The leave-one-out results are given in Table 2 beside the results for the true data. Figure 2 shows the precision-recall curve averaged over 30 simulated datasets. As in the true data analysis, the results demonstrate high precision and recall, expecially in the “nucleus” and “cytoplasm”. The “membrane” shows the worst precision-recall because we have the fewest informative data types there, but when simulating from the true data generating process, we still do quite well.

## 4 Discussion

The proposed methodology achieves fairly strong predictive power by integrating data in a compartment specific way. Importantly, we are able to evaluate how each data type contributes to the overall likelihood of any edge. Since each data type independently contributes to the probability of an edge, we can compute the fraction of the overall likelihood difference (between an edge and no edge) that is due to a particular data type. In this way our framework provides information about which parts of a pathway hypothesis are not well supported by available data (see Figure 3).

**Figure 3:**
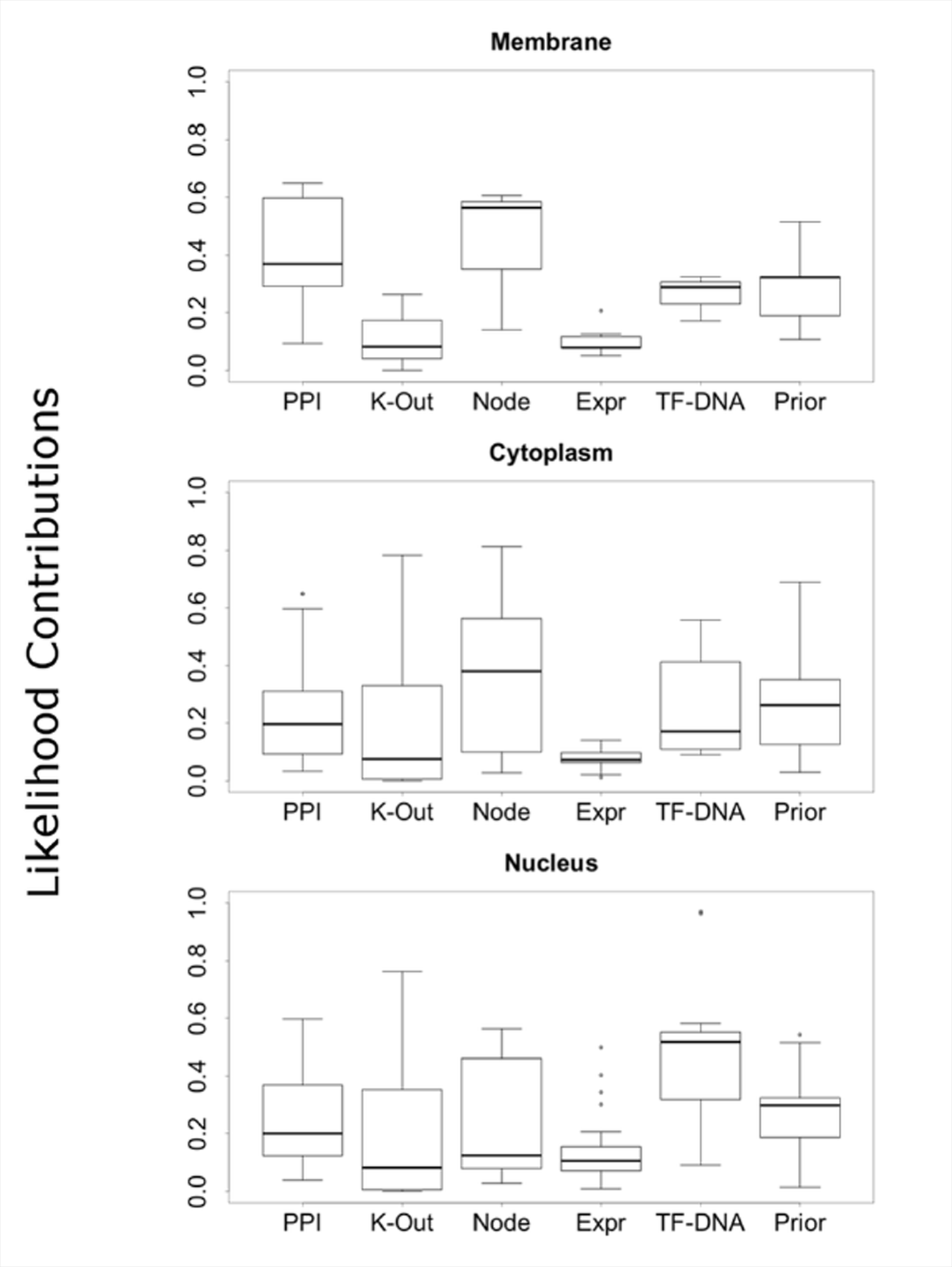
Percentages of differential likelihood (presence vs. absence of an edge) due to specific data types, by compartment. Node data contribute the most in the cytoplasm (center), whereas TF-DNA binding data contribute the most in the nucleus (right).

In addition, our methodology can identify if a particular data type tends to disagree with the other data types for sets of edges. This could indicate whether or not a data type is at all useful for modeling edges in a particular cellular location. Thus, it may be possible to do inference on the compartment map from Table 1, rather than fix it a priori. Alternatively, this information can be used to check the validity of the individual data models of Section 2.

There are some open statistical issues that could be addressed in future work. One problem with the node data, is that the protein domains are diverse and sparse. While there is evidence of signal here, there is an over-fitting problem. With more domain data, or perhaps broader domain categories, we may be able to learn more from the prior pathway. If this was the case, the leave-one-out results in the cytoplasm might improve significantly. This is evident from our results which show how borrowing domain information from other MAPK sub-pathways significantly improved the posterior probabilities of edges in the leave-one-out simulations.

We also noticed that most of the knockouts in the gene perturbation data set we used were generally downstream. If the knockouts were further upstream from perturbed genes in the nucleus, then we could learn about the possible presence of edges in a path between the knockout and other genes.

Lastly, we divided the pathway into its three main compartments: membrane, cytoplasm and nucleus. However, in future work, we hope to divide the pathway more finely into the over two dozen cellular components specified by the gene ontology (GO) for the yeast *S. Cerevisae.* By dividing the pathway into more compartments, we would also have a greater degree of control over which data types are used in various parts of the cell.

### 4.1 Concluding remarks

In this paper we introduced a technique for refining cellular pathway models by integrating heterogeneous data sources in a compartment specific way and explicitly included node properties in our model. Our case-study results indicate that this model can be useful for discovering new components or cross-talk with other pathways. Our powerful and flexible pathway modeling framework can be easily extended and modified to include additional and novel datasets.

### A Supplementary results

In this appendix we present more details about the simulation results.

**Figure 4:**
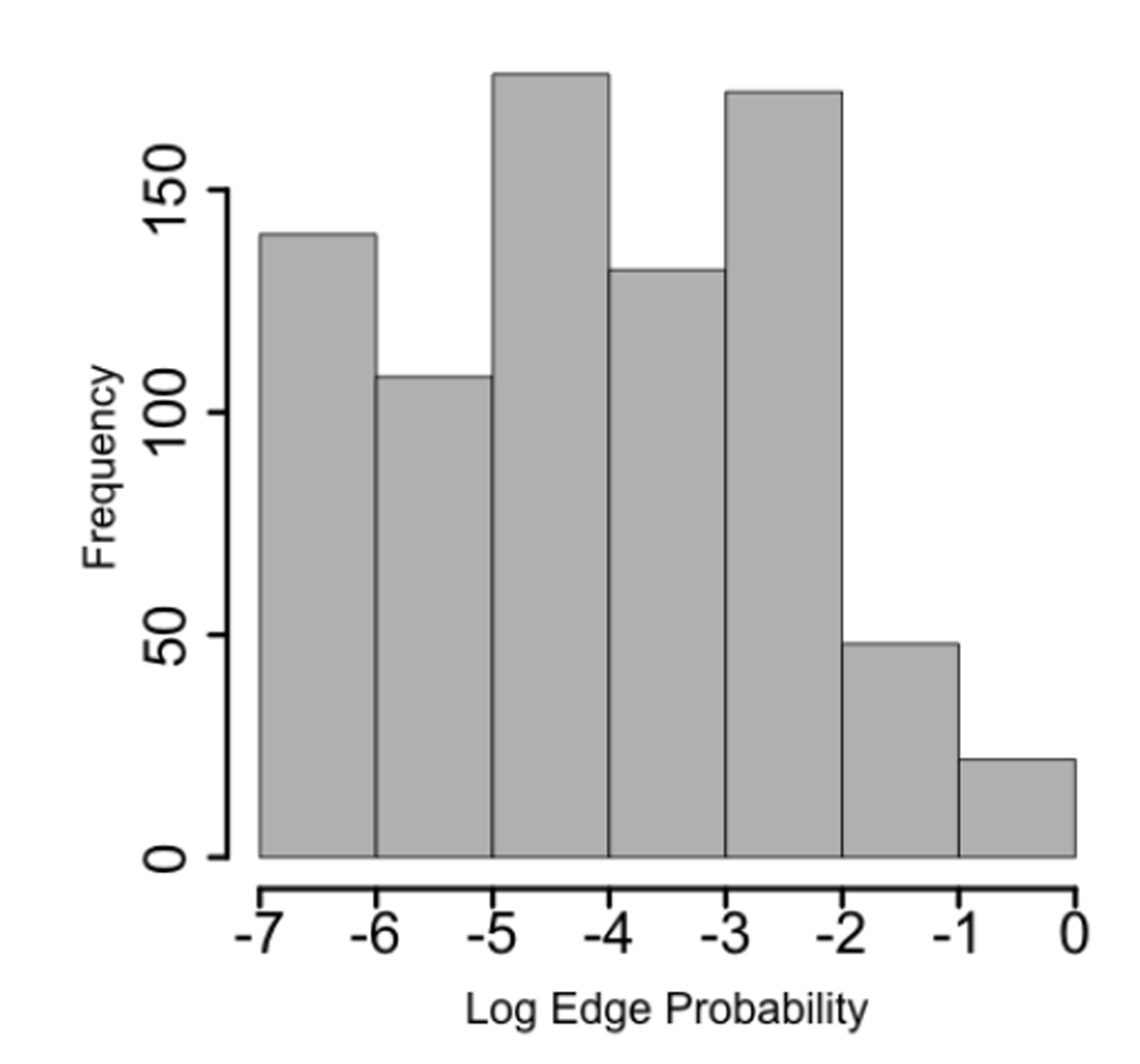
Log posterior probabilities for edges that were not in the hypothesis pathway. The vast majority of non-edges have small posterior probability (third quantile at 0.02). However, there are a few highly probable edges, which may indicate previously undiscovered interactions.

